# Rapid learning of temporal dependencies at multiple timescales

**DOI:** 10.1101/2024.01.15.575748

**Authors:** Cybelle M. Smith, Sharon L. Thompson-Schill, Anna C. Schapiro

## Abstract

Our environment contains temporal information unfolding simultaneously at multiple timescales. How do we learn and represent these dynamic and overlapping information streams? We investigated these processes in a statistical learning paradigm with simultaneous short and long timescale contingencies. Human participants (N=96) played a game where they learned to quickly click on a target image when it appeared in one of 9 locations, in 8 different contexts. Across contexts, we manipulated the order of target locations: at a short timescale, the order of pairs of sequential locations in which the target appeared; at a longer timescale, the set of locations that appeared in the first vs. second half of the game. Participants periodically predicted the upcoming target location, and later performed similarity judgements comparing the games based on their order properties. Participants showed context dependent sensitivity to order information at both short and long timescales, with evidence of stronger learning for short timescales. We modeled the learning paradigm using a gated recurrent network trained to make immediate predictions, which demonstrated multilevel learning timecourses and patterns of sensitivity to the similarity structure of the games that mirrored human participants. The model grouped games with matching rule structure and dissociated games based on low-level order information more so than high-level order information. The work shows how humans and models can rapidly and concurrently acquire order information at different timescales.

## Introduction

The environment contains temporal dependencies that play out simultaneously at multiple timescales. For example, a baseball fan can anticipate the trajectory of a just-pitched ball as well as the switching of jerseys in the field between the top and bottom half of each inning. To fully represent, simulate, and anticipate changes in the environment, humans must learn temporal dependencies that span multiple timescales. Such learning across timescales has been documented in a variety of domains, including language (Saffran & Wilson, 2003), event segmentation (Davachi & DuBrow, 2015; Shin & DuBrow, 2021), motor learning (Krakauer et al., 2019), and visual statistical learning (Schapiro et al., 2013; Karuza et al., 2017). Our study adds to this literature by testing whether humans can rapidly learn simultaneous regularities occurring at multiple timescales. In particular, we explore how sensitive humans as well as neural network models are to slow background statistical dependencies when they are focused on making short-term predictions in the immediate task at hand.

Studies on statistical learning of auditory content (language and music) suggest that multiple timescales of temporal statistics can be rapidly acquired under at least some circumstances. For example, infants can learn transition probabilities among both syllables (Saffran et al., 1996) and pairs of syllables (“words”; Saffran & Wilson, 2003) after only a few minutes of exposure. Moreover, infants and adults are sensitive to non-adjacent dependencies among nonsense words within a single learning session (Gómez, 2002; Misyak et al., 2010), and adults can learn higher order statistics among tones after a mere five minutes of exposure (Furl et al., 2011). In these cases, higher order learning is often modulated by the salience of embedded fast temporal statistics. For example, the nature of intervening sounds can impact learning success for long distance dependencies. In particular, the presence of high adjacent transition probabilities impedes acquisition of non-adjacent dependencies (Gómez, 2002), raising the possibility of attentional trade-offs for statistical learning at different temporal scales. Indeed, when intervening sounds are of a different type from those related through a “non-adjacent” dependency, the dependency becomes easier to learn (Creel et al., 2004; Newport & Aslin, 2004). It is unknown whether these salient differences are needed when the higher order dependencies unfold at a slower timescale (more than a few seconds), where they may be less likely to be confused with the short timescale information.

In the visual, spatial, and motor domains, humans rapidly learn short temporal dependencies across a variety of paradigms in under an hour of exposure (Fiser & Aslin, 2002; Krakauer et al., 2019). However, many paradigms showing sensitivity to longer timescale statistics appear to involve extensive training. For example, given many sessions of training, participants in visuospatial search and visuomotor learning tasks appear to be able to use temporal context going back at least three trials to anticipate the upcoming target location (Cleeremans & McClelland, 1991; Lewicki et al., 1987). Similarly, in the motor learning literature, after extended training, shuffling previously learned motor chunks results in a decrement in performance relative to intact sequences, consistent with learning of higher-level order among the chunks (Sakai et al., 2003). It is unclear whether there are contexts in which simple but slow visuospatial or motor statistics can be acquired rapidly, and whether the presence of faster temporal dependencies interferes with learning of slower visuospatial or motor regularities.

We report the results of a preregistered study in which humans learned nested temporal dependencies at fast and slow timescales in a visuo-spatial motor learning task (cf. the carnival game known as “whack-a-mole”). Participants played different “mini-games” of “whack-a-mole” in which the temporal dependencies among target locations on different trials varied. Participants were periodically asked to predict the upcoming target location. They were also asked to judge the similarity of the different mini-games on the basis of their temporal structure (both fast and slow). Using the results from both the prediction task and the similarity judgement task, we were able to track participant learning over time and assess whether both slow and fast timescale dependencies were being learned within a single session. We predicted on the basis of our pilot data that participants would be able to learn both slow and fast timescale dependencies, and that their knowledge would be reflected in both tasks. We also predicted that fast timescale dependencies would be learned better than slow timescale dependencies.

We simulated performance on our behavioral tasks (online predictions and post-training similarity judgements) using a gated recurrent neural network with a single hidden layer. We trained the model to predict the subsequent target location, using input stimuli generated in the same way as our human behavioral experiment, and evaluated to what extent the model became sensitive to longer timescale dependencies over training, and its degree of match to the human behavioral data.

## Methods

### Human Behavioral Methods

#### Subjects

103 participants were recruited for course credit or monetary compensation from the University of Pennsylvania participant pool. We obtained informed consent from all participants in accordance with the University of Pennsylvania IRB. Participants were native English speakers with normal/corrected-to-normal vision and hearing. We used preregistered exclusion criteria as follows: 1) Similarity judgement task attention check accuracy below 70%, 2) During gameplay, missing more than 25% of responses, 3) During gameplay, not responding on any trial for two consecutive games or more. Following these criteria, 7 participants were excluded and replaced, leaving our final pre-determined sample size of 96 participants (aged 18 to 59, mean = 25.2, 14 participants were over age 30, 5 participants did not disclose their age). Our target sample size was determined by power analysis of a smaller pilot experiment (N=32) using measures derived from the similarity judgement task described further below, with the simr package powerSim, indicating that power was over 95% for all key contrasts (without correction for multiple comparisons). Preregistered methods are available at https://archive.org/details/osf-registrations-egukx-v1.

#### Materials

Participants played 8 mini-games of “whack-a-mole.” Each mini-game contained a distinctive background image (similar to a game board or arena) and a thematically related target image (always an animate object; Fig. 1). Both the background and target images were in cartoon style. Background images were all covered with gridlines of similar size, and a gray semi-transparent overlay was used to delineate the arena during gameplay. The following background and target images were used for the 8 minigames: 1) Background: desert with skulls carved into a plateau, Target: skull; 2) Background: forest temple, Target: fairy; 3) Background: desert with pyramid structure and pond, Target: camel; 4) Background: forest with blossoming cherry trees, Target: animate cherry blossom with face; 5) Background: snowy temple with bridge and large statues, Target: yeti; 6) Background: terraced land with large mushrooms, Target: animate mushroom with face; 7) Background: underground river and rock formation with crystals, Target: bat; 8) Background: medieval encampment, Target: dragon. The order of locations in which the target image appeared in each game was counter-balanced across participants, such that no background/target image pair was consistently associated with any set of order rules. Participants were given an additional practice game in which the target appeared in completely random locations. The background and target images used for this game were distinct from those used during training, and consisted of a swimming pool and a goldfish.

**Figure 1.**
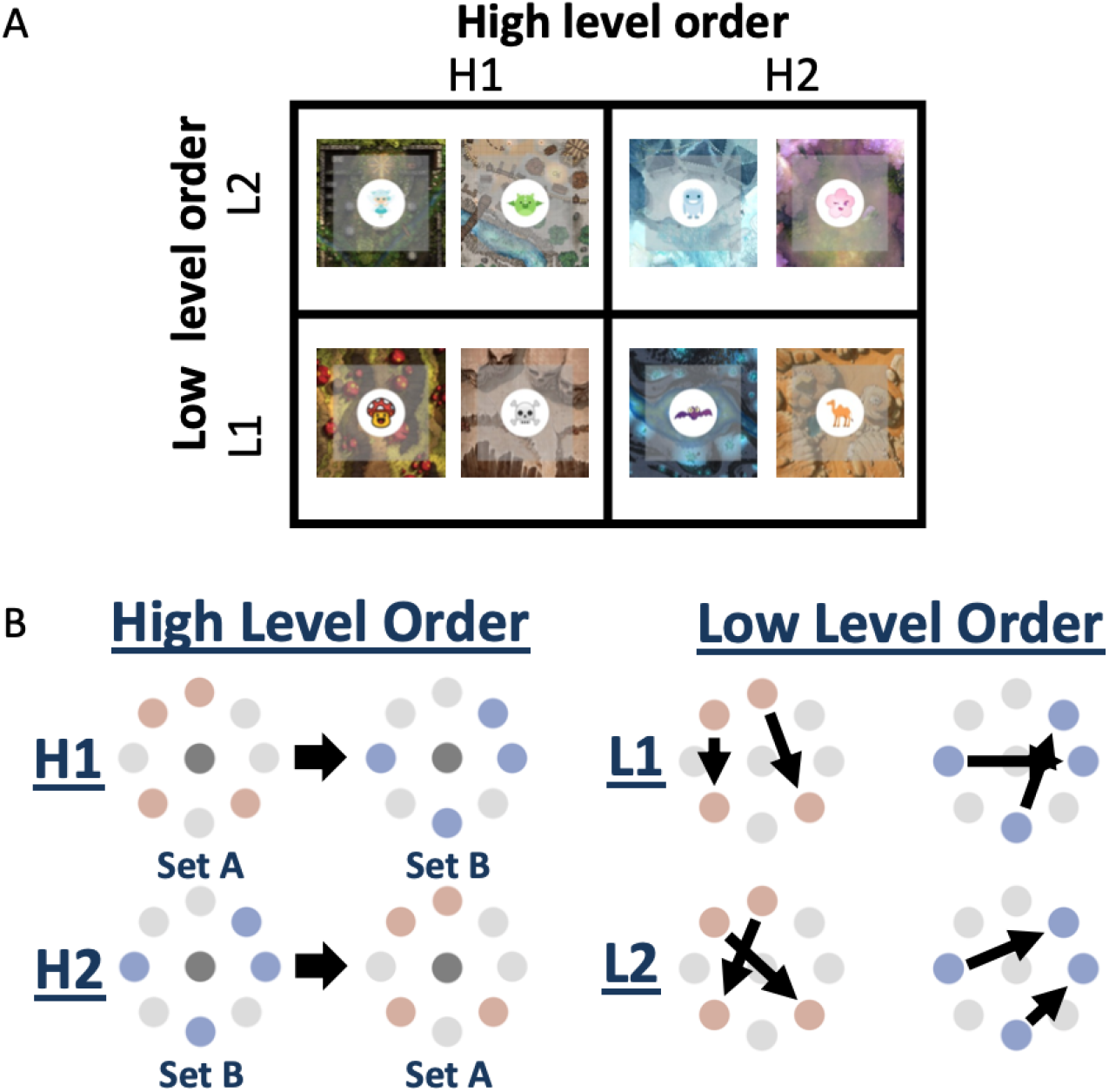
Experimental conditions. (A) High and low level order rules are manipulated within subjects in a 2×2 design. (B) High level order determines the set of target locations that is encountered in the first vs. second half of the game. Low level order determines which paired associations are encountered among the target locations on adjacent trials during the game.

## Procedure

### Exposure to games

Participants were informed that they would play a series of mini-games similar to “whack-a-mole.” They were asked to pay attention to how similar the mini-games were to each other and informed that they would answer questions comparing the games afterwards. They were particularly asked to pay attention to the order in which the target appeared during the games, and differences in that order among the games. Participants played each of 8 mini-games once per round in a series of 8 rounds. Participants received points for their performance and could view the point total for each game during that game, and their total point total after each game. Each mini-game began with a slide that introduced the background of the game (e.g., a drawing of a snowy mountain temple scene with gridlines over it), and the target image that they needed to click (e.g., a cartoon yeti), which was superimposed on the background. During each mini-game, a board was displayed with 9 dark circles overlaid on it that resembled “holes”: one center hole and 8 holes evenly spaced on the periphery in a circular arrangement. During each trial of the game, a target image appeared in one of the 9 holes and the participant had to click on the target before it disappeared to receive 25 points. The time limit to receive points was 800 ms after target onset. If they did not click the target in time, they heard a laughing noise and did not receive any points. Participants were informed prior to playing the games that the laughing noise indicated that they were too slow. Clicking the target ended the trial. If the participant did not click the location that the target appeared in, the trial ended 2200 ms after target offset. The intertrial interval was 500 ms (from end of one trial to appearance of target in the next trial).

### Game structure

Each game lasted 27 trials. Games were divided in a 2×2 design by adopting one of two high level order rules (slow timescale) and one of two low level order rules (fast timescale; Figure 1). Two of the 8 games were assigned to each combination of high and low level order rules. The 8 peripheral (non-center) target locations were randomly partitioned into 2 sets of 4 peripheral locations. High level order rules governed which set (A or B) was used for target locations in the first half of the game; in the second half of the game the target appeared in the remaining set of locations. Locations were also grouped into ordered pairs such that if one peripheral location was visited on a given trial, the target would appear in the second location on the following trial 100% of the time. Low-level order rules governed the second location in each of the four ordered pairs of locations. The second location of each pair was swapped across games with different low-level order. Prior to presenting each ordered pair of locations, the target appeared in the center. In the middle of each game, the target appeared in the center three times in a row to help form an event boundary separating the first and second half of the game.

All participants were assigned to a unique randomized list of games, belonging to one of 16 counterbalancing groups. Counterbalancing groups were designed such that each game stimulus was assigned to each of the four joint order conditions (H1L1, H1L2, H2L1, H2L2) an equal number of times across lists. Each game stimulus shared its high level or low level order condition with each other game stimulus an equal number of times across the counterbalancing. The order of presentation of paired associate locations within each half of the game was randomized. Two pairs were presented twice in each half of gameplay (the remaining two pairs were presented twice in the second half). The games were played in “rounds” of 8 such that each mini-game was played once per round. The order of the games within a round was randomized.

### Prediction probe trials

During game-play participants were periodically probed on their order knowledge by explicitly predicting the upcoming target location. During probe trials the 8 peripheral locations were covered by a “?” and the participant had to select where they thought the target would appear next (barring the center hole). Responses were self-paced. Each probe trial was followed by a 1000 ms delay and then by an ordinary game trial that provided participants with feedback on their predictions. There was one prediction probe trial each time a game was played, except for the final round, when there were three probe trials per game. Prediction probe trials occurred throughout the game at all non-center positions but were more likely to appear as the second trial in the game (37.5% of the time, then roughly evenly distributed across remaining trial numbers and positions). Participants received 750 bonus points each time they selected the actual target location. Depending on when the trial appeared within the game, maximum possible accuracy was either 50% (for the first location in an ordered pair) or 100% (for the second location in an ordered pair).

### Similarity judgement task

After playing the games, participants judged the similarity of the games in a two-alternative forced choice task. Participants were presented with a picture representing one game at the top and were asked which of two games presented at the bottom was more similar to the game at the top. Figure 1A displays the full set of stimuli presented to represent the games. All participants were asked to base their judgements on the order of locations in the games. They were randomly assigned to one of two conditions for finer grained instructional wording (48 participants per condition): in condition 1, they were told to base similarity on “when/where the target appeared”, and in condition 2, on the “sequence of locations [the] target appeared in.”

Similarity judgements were performed on distinct trial types:

a. Attention check. One choice was the same game as the comparison game (with the same target image). This trial type was used for performance-based exclusion.
b. Same rules (but different game identity / target image) vs. game with different high and low level order condition. Performance on this trial type may reflect sensitivity to either low or high-level order information.
c. Same rules vs. game with different low level order condition. Above chance accuracy would reflect sensitivity to low-level order information.
d. Same rules vs. game with different high level order condition. Above chance accuracy would reflect sensitivity to high-level order information.
e. Both low and high level order differ vs. only high level order differs from the comparison game. Above chance accuracy would reflect sensitivity to low-level order information.
f. Both low and high level order differ vs. only low level order differs from the comparison game. Above chance accuracy would reflect sensitivity to high-level order information.
g. One game matches the comparison game on low level order, and the other matches the comparison game on high level order. This trial type assesses bias towards relying on high vs. low level order information when making similarity judgements.

## Analysis

Logistic regression models were fit with the glmer function using the lme4 package in R. Nested model comparisons with the anova function were used to test for significant effects. The emmeans package was used for statistical reporting of condition means. Reported p-values are uncorrected except where specified.

### Online prediction judgements

Following our preregistered analysis plan, we fit logistic regression models to assess indices of sensitivity to high and low level order information and their change over time. Low-level sensitivity was assessed using responses to probes at the second location among paired locations. It was computed as: [Proportion congruent with low-level order rule] – [Proportion congruent with opposite low-level order rule]. High-level sensitivity was assessed using responses to probes at the first location among paired locations. It was computed as: [Proportion congruent with high-level order rule] – [Proportion congruent with opposite high-level order rule]. This was equivalent to subtracting the proportion of trials with responses drawn from the incorrect set of four locations, given the game rules and game half, from the correct set of four locations. High-level sensitivity measures were also computed separately using only the first non-center trial of each game. Responses to the first non-center trial in each game indexed sensitivity of location predictions to the target and context image. Our full model used the following formula in R notation:

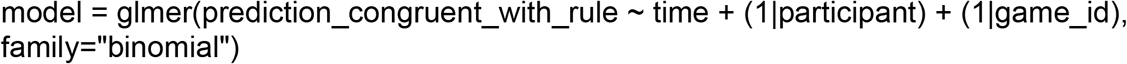

Here, time was indexed as the number of individual 27-trial games played since the beginning of the session, divided by 100 to aid with parameter estimation. We included crossed random intercepts for participant and game identity (which of the 8 games was being played, in terms of the target and context image; since game positions were assigned to random locations for each unique participant, there was no reason to anticipate consistent effects of rule condition across participants). Separate models were fit for modeling sensitivity to the high and low level rules. Reduced models excluding time as a variable were used to assess mean performance across the entire session. Nested model comparisons were used to assess the presence of an interaction between sensitivity and time.

### Similarity judgements

We fit a logistic regression model to the similarity judgement data, including trials that tested high-level order knowledge, low-level order knowledge, or both, with the following formula in R notation:

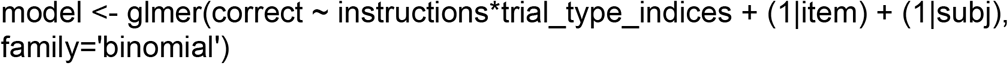

The variable trial_type_indices was a factor with 3 levels coding whether a trial type was an index of sensitivity to high-level order, sensitivity to low-level order, or sensitivity to both; thus the high and low-level order levels were each aggregated across two different original trial types, in which one option matched the target on either both or neither dimension. We tested for the significance of fixed effects of interest using the emmeans package and nested model comparisons. We also ran a second logistic regression model and evaluated its output in the same way, but with disaggregated trial types (i.e. a 5-factor model of trial type with a factor for each of trial types b-f above). We also separately examined performance on trial types a (attention check) and g (pitting high and low-level order information against each other) using t-tests.

## Modeling Methods

### Architecture

A Gated Recurrent Unit (GRU) model with a single hidden layer (N=48 model instances) was trained to predict the upcoming location in a whack-a-mole task with identical statistical properties to the human participants. We chose this architecture for its ability to effectively learn long timescale dependencies (Shewalkar et al., 2019). Like humans, the model was probed on its performance on location prediction and judgements of similarity among the games. The model had a 1x17 input layer, a 1x150 layer of gated recurrent hidden units (Chung et al., 2014), and a 1x9 output layer (Fig. 4a). All layers were fully connected. The 1x17 input vector consisted of 8 context units denoting the one-hot encoded game ID and 9 one-hot encoded location units (representing one center and 8 peripheral target locations). Gated recurrent units learn weights to generate a candidate hidden activation based on their input and their own previous activation, and then separately learn how much to integrate the candidate hidden activation with the previous activation (again based on both the input and the previous activation) to generate their output. The model was implemented in Python 3.6.10 using keras tensorflow (tensorflow v. 2.1) with default fitting parameters and a Mean Squared Error loss function in batches of 8 games at a time (equivalent to a single round of exposure to all 8 games). Activations were reset in between games. The effective learning rate was adjusted automatically over training using the RMSprop optimizer to speed convergence.

### Training

Models were trained on sequences generated using the same randomization and counterbalancing procedures as human participants, for 240 rounds of games. Preliminary examination suggested that the model performed similarly to human participants on prediction and similarity judgement tasks after receiving approximately 5 times the exposure that human participants did (40 rounds of games), given a default global learning_rate parameter of 0.001. Presented analyses probe the model after 5x human exposure, unless otherwise specified. Separate model instances were initialized with random weights as follows: connection weights between layers were taken from a Glorot uniform distribution (Glorot & Bengio, 2010), recurrent weights were computed from an orthogonal matrix derived from an initial random normal matrix, and bias weights were initially set to zero.

### Analysis

#### Online predictions

At various stages of training, weights were frozen and the model was asked to predict the next location at the same probe points as the human participants on a number of trials (172800 trials over 800 rounds of the 8 games). Max activation of the output units was used to determine the most likely predicted location. The same sensitivity measures were derived as for humans: low-level sensitivity, and high-level sensitivity at the earliest probe point in each game. Gaussian noise was applied to all 17 input units prior to making a prediction. Several noise levels were assessed (0.01, 10, and 0.1-2.0 in increments of 0.1) and a Gaussian with a standard deviation of 1.0 was determined to best correspond to human data. To create stable performance estimates, performance was averaged over 10 injections of noise.

#### Similarity judgements

After training, model weights were frozen and the model was exposed to inputs such that a single game ID unit was turned on and all other input units were off. Activation of the model’s hidden units was compared across games using Pearson correlations as a metric of distance. The model was then asked to compare games’ similarity using trials generated in the same way as for human judgements. A softmax function was used to determine the probability of each choice based on its relative distance to target (temperature = 1), and we sampled from the probability distribution ten times for each trial to generate probabilistic choice data, then took the average performance across trials for each trial type and model instance. Trial types were analyzed analogously to the human data.

#### Activation trajectories

Hidden unit activations were recorded during game play. The average hidden activation vector was taken across all rounds for each game ID and trial (from 1-27 during the game) for each model instance. Then distance matrices between all the averaged activity vectors were computed for each model instance, and the distance matrices were averaged across model instances. Multidimensional scaling (in two dimensions) was applied to the final aggregate distance matrix using the sklearn.manifold.MDS function from the scikit-learn package (version 0.24.2) in python (version 3.6.10) with default parameters, and the resulting mean activation trajectories over the course of game play were plotted.

## Results

### Human Behavior

#### Online predictions

Following our preregistered analysis plan, we fit logistic regression models to the online prediction data. In our assessment of high-level sensitivity, participants were above chance at predicting locations from the correct half of the game (earliest probe point: mean = .266, SE = 0.060, Z ratio = 4.424, p < .001; all probe points: mean = .525, SE = .053, Z ratio = 9.891, p < .001) and did so more often later in training, although not significantly so when only the earliest probe point was considered (earliest probe point: β = .310, SE = .195, 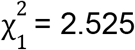, n.s.; all probe points: β = .410, SE = .150,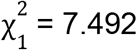, p= .006). Participants were also more likely to select the appropriate second location in a pair than the location corresponding to the opposite low-level order rule (β = .892, SE = .112, Z ratio = 7.953, p < .001). Again, this trend increased over the course of training (β = 1.605, SE = .326, 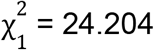, p < .001).

We also report t-tests corresponding with Figure 2. In the second half of training, humans were above chance on our measures of both low and high level sensitivity (low level sensitivity: mean = .298, t(95) = 8.073, p < .001; high level sensitivity (all trials): mean = .264, t(95) = 9.305, p < .001; high level sensitivity (earliest trial in game): mean = .142, t = 4.222, p < .001). High level performance was higher measured across all trials than at the earliest trial in game (difference = .122, t(95) = 5.550, p < .001). Low level performance did not differ from high level performance across all trials (difference = -.034, t = -.947, n.s.). However, low level performance (across all trials) was higher than high level performance at the earliest trial in game (difference = .155, t(95) = 3.295, p = .001).

**Figure 2.**
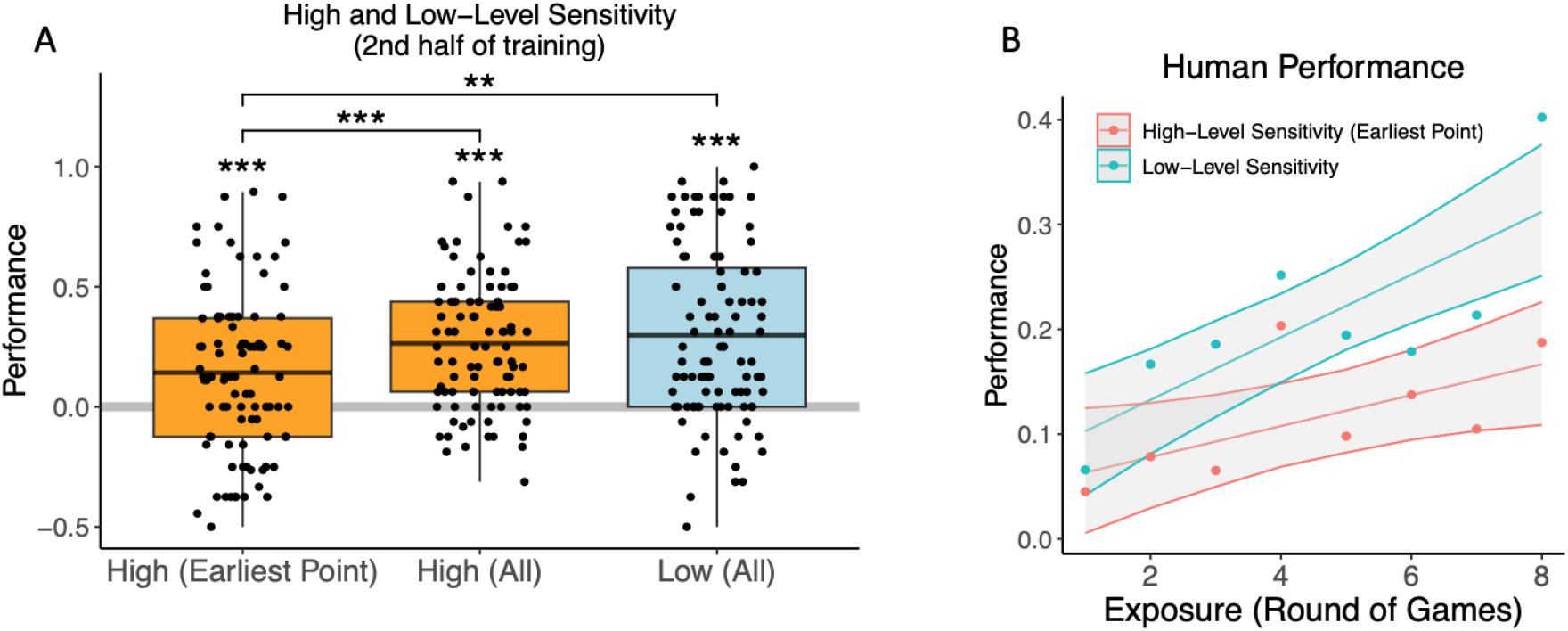
Online performance measures of learning in human participants. (A) Participants demonstrated sensitivity to high and low level order rules in their online predictions of upcoming locations during the second half of training. Boxplot midlines show the mean and error bars indicate upper and lower quartiles, ** p < .01, *** p < .001. (B) Sensitivity to high and low level order information plotted by round of 8 games over the course of training, shaded regions indicate bootstrapped 95% confidence bands.

Overall, participants learned to predict locations in a manner indicating sensitivity to both low and high level structure, with some evidence for stronger low level sensitivity.

#### Similarity judgements

Performance was marginally better in the similarity judgment task when participants were asked to base their judgements on “when/where” the target appeared, rather than the “sequence of locations” that the target appeared in (3 trial type scheme: β = -0.255, SE = .134, 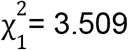, p = .061). Adding an interaction term between instruction χcondition and 3-way trial type – high, low or both – did not improve model fit (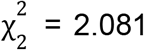, n.s). We specifically predicted that instructions condition would modulate sensitivity to low-level order on the similarity judgement task, and this was not born out (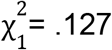, n.s.). Thus, we collapsed data across instruction condition for subsequent analyses.

Using logistic regression as per our preregistered analysis plan, but now collapsing across instructions condition, human participants were above chance on trials indexing sensitivity to low-level order and to both low and high-level order combined (low: β = .207, SE = .053, Z ratio = 3.938, p < .001; both: β = .278, SE = .068, Z ratio = 4.096, p < .001). They were only marginally above chance on trials indexing sensitivity to high-level order (β = .088, SE = .052, Z ratio = 1.674, p = .094). Participants had higher accuracy on trials that probed low-level sensitivity than on trials that probed high-level sensitivity only (high vs. low odds ratio = .888, SE = 0.0378, Z ratio = -2.793, adjusted p = 0.015; both vs. high odds ratio = 1.209, SE = .0732, Z ratio = 3.142, adjusted p = .005; p-values adjusted by the Tukey method). Trials that probed low-level sensitivity did not differ in accuracy from trials reflecting both high and low level sensitivity (both vs. low odds ratio = 1.074, SE = .065, Z ratio = 1.177, n.s.).

Broken down further by trial type (5-way scheme), participants were above chance on all trial types except for one measure of high-level sensitivity (“two different rules vs. same low-level rule”): (b) both same rules vs both different rules, est. mean probability correct = .569, SE = .0166, Z ratio = 4.094, p < .001; (c) same rules vs low level different, est. mean probability correct = .563, SE = .0167, Z ratio = 3.718, p < .001; (d) same rules vs high level different, est. mean probability correct = .535, SE = .0168, Z ratio = 2.074, p = .038; (e) both different rules vs only high level differs, est. mean probability correct = .546, SE = .014, Z ratio = 3.250, p = .001; (f) both different rules vs only low level differs, est. mean probability correct = .515, SE = .0141, Z ratio = 1.088, n.s.

Humans showed above chance accuracy on the attention check (mean = .970, t(95) = 77.125, p < .001) and were significantly biased towards using low level order information when high and low level information sources were directly pitted against each other (trial type g; mean =.458, t = -2.543, p = .013).

T-tests corresponding with Figure 3 are reported below for each trial type:

**Figure 3.**
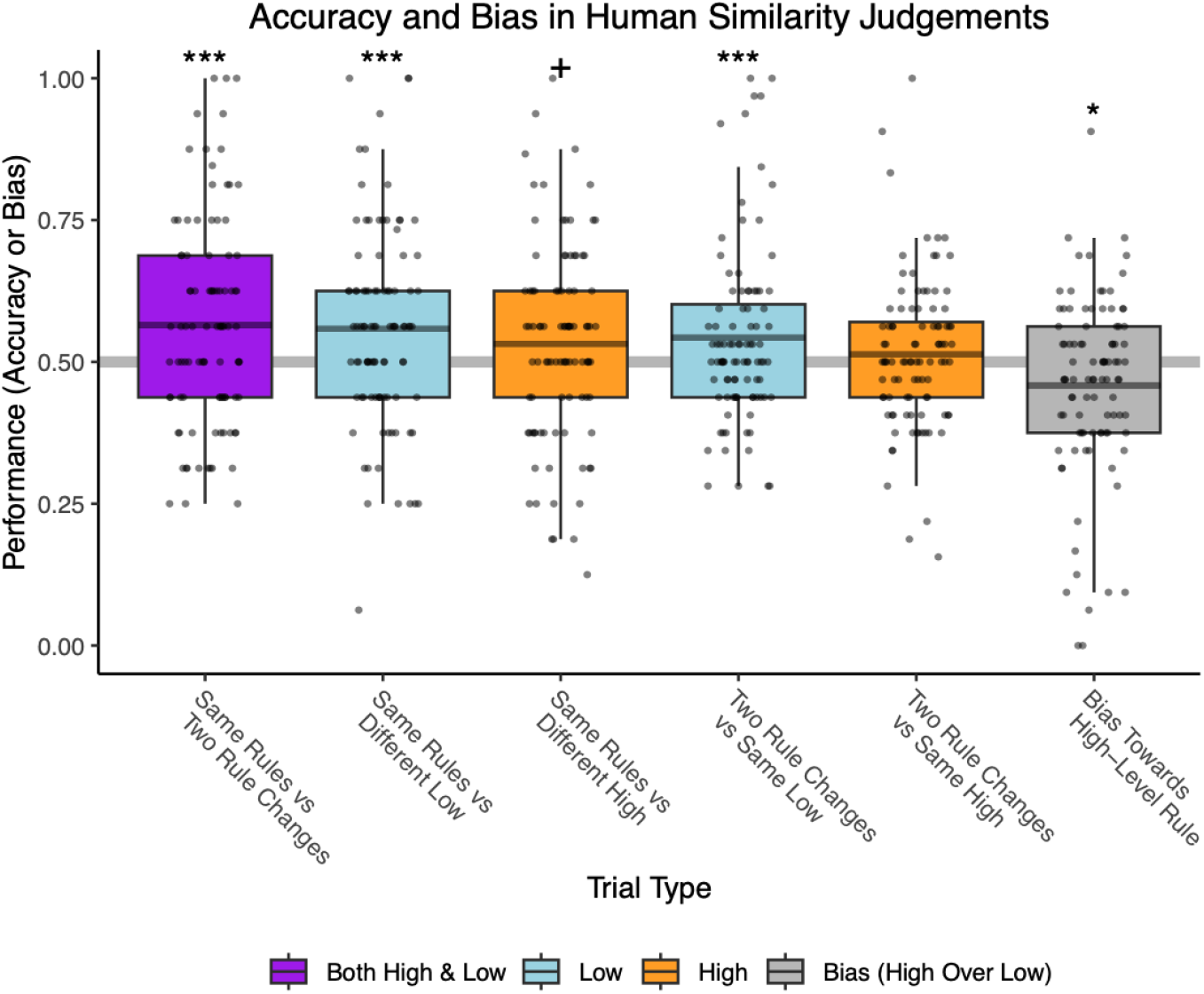
Human similarity judgements. Participants were more sensitive to low than high level order information in their similarity judgements. Boxplot midlines show the means, + p < .1, * p < .05, *** p < .001.

**Figure 4.**
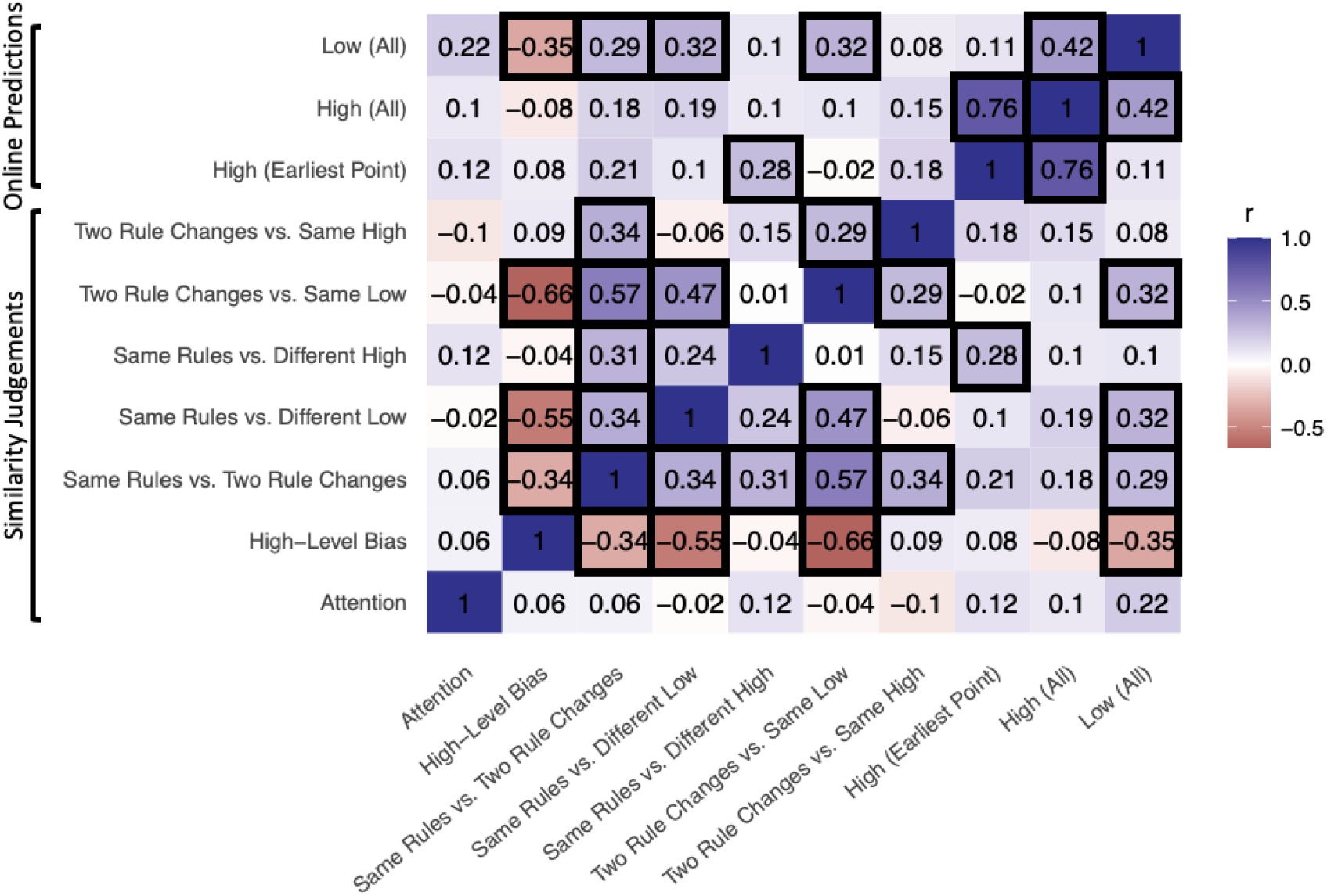
Correlations among human behavioral measures. Pearson correlations surviving FDR correction (q < .05) outlined in black.

b) Same rules (but different game identity / target image) vs. game with different high and low level order condition. Mean = .564, t(95) = 3.451, p < .001.

c) Same rules vs. game with different low level order condition. Mean = .558, t(95)= 3.268, p = .002.

d) Same rules vs. game with different high level order condition. Mean = .532, t(95) = 1.804, p = .074.

e) Both low and high level order differ vs. only high level order differs from the comparison game. Mean = .543, t(95) = 2.678, p = .009.

f) Both low and high level order differ vs. only low level order differs from the comparison game. Above chance accuracy would reflect sensitivity to high-level order information. Mean = .513, t(95) = .978, n.s.

Overall, both high and low level order influenced participants’ similarity judgments, but judgments were more strongly impacted by low level order information.

#### Correlation analysis

We further examined by-subject correlations among sensitivity measures from our online prediction and similarity judgement tasks (Figure 4). We tested all possible correlations between all of the above reported behavioral measures across both tasks, and corrected for the False Discovery Rate (FDR) using Benjamini and Hochberg’s (1995) procedure (q = 0.05; reported p values and confidence intervals are uncorrected, but all significant results survived FDR correction). In this way, we found that measures of sensitivity to low-level order information tended to be correlated with each other across tasks, and the same was also true for high-level order information. Specifically, sensitivity to low-level order on the online prediction task was correlated with both measures of low-level sensitivity on the similarity judgement task (“same rules vs. different low”: r = .321, 95% CI = [.128, .490], t = 3.281, p = .001; “two rule changes vs. same low”: r = .319, 95% CI = [.127, .488], t = 3.264, p = .002). Sensitivity to high-level order on early trials in game play on the online prediction task was correlated with high-level sensitivity on “same rules vs. different high” trials in the similarity judgement task (r = .280, 95% CI = [.084, .455], t = 2.824, p = .006). While there was a significant positive correlation between one measure of online sensitivity to high-level order and online sensitivity to low-level order (r = .422, 95% CI = [.242, .574], t = 4.517, p < .001), the purer measure of high-level sensitivity (at the earliest points in game play) did not correlate with low-level order information (the measures in Fig. 2B; r = .106, 95% CI: [-.096,.300], Bayes Factor against r=.333 was .388), suggesting some degree of decoupling of these measures.

**Figure 5.**
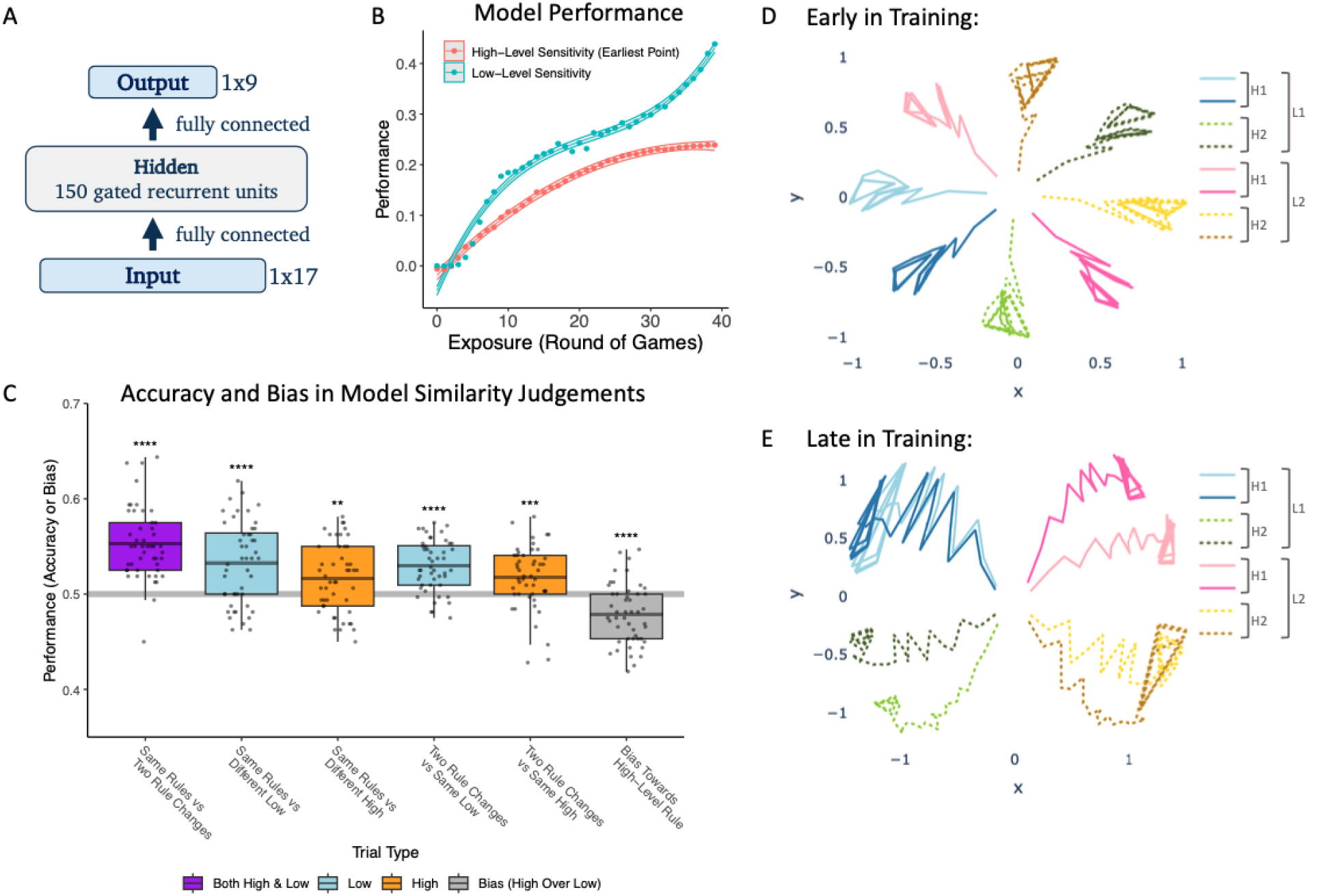
Modeling results. (A) Model architecture. (B) Average model performance on online prediction task over training. Shaded regions indicate bootstrapped 95% confidence bands. (C) Model similarity judgements after 40 rounds of training. Boxplot midlines show the means, ** p < .01, *** p < .001, **** p < .0001. (D and E) Mean hidden activation trajectories during the course of game play for each of the 8 games early in training (D, 8 rounds), and late in training (E, 40 rounds). Hidden activation trajectories start out roughly equidistant among the 8 games but become clustered by high and low level order rules over the course of training.

#### Neural network model

##### Online predictions

After 40 epochs of training (exposure to 320 games), the model showed above chance low-level sensitivity (mean = .400, t(47) = 31.145, p < .001) and high-level sensitivity (mean = .239, t(47) = 85.008, p < .001) in its predictions. Like human participants, the model demonstrated higher performance on our measure of low-level sensitivity than on high-level sensitivity over the course of learning (Figure 4b; difference by 40 rounds exposure = .161, t(47) = 11.479, p < .001).

##### Similarity judgements

The model also showed a pattern of similarity judgements that was qualitatively similar to human participants (Figure 4c). Individual tests of above-chance accuracy and high vs. low level bias are reported below for each trial type:

a. Same rules (but different game identity / target image) vs. game with different high and low level order condition. Mean = .553, t(47) = 9.682, p < .001.
b. Same rules vs. game with different low level order condition. Mean = .533, t(47) = 5.273, p < .001.
c. Same rules vs. game with different high level order condition. Mean = .516, t(47) = 3.143, p = .003.
d. Both low and high level order differ vs. only high level order differs from the comparison game. Above chance accuracy would reflect sensitivity to low-level order information. Mean = .530, t(47) = 7.759, p < .001.
e. Both low and high level order differ vs. only low level order differs from the comparison game. Above chance accuracy would reflect sensitivity to high-level order information. Mean = .518, t(47) = 3.661, p = .001.
f. High-level bias. One game matches the comparison game on low level order, and the other matches the comparison game on high level order. Mean = .479, t(47) = -4.645, p < .001.

Sensitivity to trials that probe low level order was higher than to those probing high level sensitivity (mean difference = .014, t (47) = 2.767, p = .008).

##### Model hidden activation trajectories

We examined the model’s hidden activation trajectories during game play for each game ID at both early (8 rounds of games; Figure 4d) and late (40 rounds; Figure 4e) stages of training. Trajectories are shown branching out from the center of the plot as game play continues from the start to end of each game. Early in training, the model represented individual games differently, and all games were approximately equally dissimilar (relative to changes over time within the game). Later in training, the model grouped games by high and low level order rule. Games that differed in their low level order rule were more distinct in activation space than games that differed in their high level order rule (mean of [[distance for different low level rule – same low level rule] - [distance for different high level rule – same high level rule]] = .396, t(47) = 6.103, p < .001). Stronger representational warping according to low level order corresponds to stronger sensitivity to low level order in the model’s similarity judgments as well as online predictions.

#### Discussion

In this study we have shown that humans can rapidly learn statistical information at slow (high-level) temporal scales while performing a task that focuses on concurrent fast (low-level) statistics. We have also captured aspects of human behavior using a recurrent neural network trained to predict immediate upcoming input, which recapitulated patterns in human online predictions and similarity judgements. Modeling results suggest that a common learning and representation scheme can capture information at multiple temporal scales. Both the human and modeling results were consistent with prioritization of learning for rapid timescale statistical relationships over slower timescale relationships, and with an account in which learning at the two levels unfolds in parallel.

Humans were able to learn context-dependent high-level order information while performing a visuo-motor task (whack-a-mole) in which they also learned low-level order information. Participants’ low-level order knowledge was demonstrated both in their online predictions during training and in their post-training similarity judgements. Their high-level order knowledge was demonstrated in their online predictions (even when based on the visual context alone) and on similarity judgement trials that pitted a match on high and low level order against a high level mismatch. It also was reflected in the positive correlation between high level order sensitivity on the online prediction task (on early trials using visual context) and the similarity judgement task. In both online predictions and similarity judgments, there was evidence for greater sensitivity to low-level statistics.

The fact that humans were able to rapidly gain sensitivity to high-level order statistics while performing a task focused on learning low-level order statistics is not trivial, since the presence of low-level dependencies has been suggested to interfere with learning of higher level dependencies in some domains (e.g. Gómez, 2002; Newport & Aslin, 2004). Our findings are consistent with faster temporal statistics being represented differently enough from these kinds of slower temporal statistics during visuomotor learning that they do not provoke interference. Also consistent with this interpretation, sensitivity to high and low level order information was not strongly correlated across subjects in our paradigm. However, dissociability in the representations of slow and fast temporal statistics is not incompatible with the idea that both could be acquired using a common learning mechanism and neural substrate, as suggested by our modeling results.

A gated recurrent neural network with a single hidden layer is able to learn statistical patterns unfolding at both slow and fast timescales, and approximates human behavior in our paradigm. The model makes online predictions that match human low and high-level sensitivity measures. The model also, like humans, demonstrates greater sensitivity to low than high level order information in the similarity judgement task. Our close match to the human data at multiple timescales is notable given that our model was only trained to predict the upcoming target location. Training our model to predict the upcoming target location could result in a close match to our human data because it matches our explicit predictive probe task, and/or because it is consistent with a broader prediction-based learning account of statistical learning (e.g., Friston, 2005; Kiebel et al., 2008; Schapiro et al., 2013). It is also interesting that our model was able to learn statistical dependencies at multiple timescales given that it uses a single, unified learning mechanism. Our model learns what temporal scales are relevant, rather than having them explicitly parameterized, and accomplishes learning with a single hidden layer. This is in contrast with models such as the Hierarchical Autoencoders in Time (HAT) model (Chien & Honey, 2020), which employs a more constrained gating mechanism that is modulated by layer depth and explicit temporal integration parameters in order to learn statistics at multiple temporal scales. Our model’s success despite its flat structure is consistent with prior demonstrations that single layer models can capture complex multilevel temporal dynamics (e.g., Botvinick, 2007; Botvinick & Plaut, 2004).

Our model also demonstrates that slow and fast timescale statistical relationships can be learned concurrently. Although the model did show bias towards enhanced learning of low-level / fast temporal statistics, it improved on sensitivity measures for both fast and slow timescale statistics simultaneously. This matches visual trends in the human data (Fig. 2B), although we were not powered to examine this directly. Simultaneous learning at multiple timescales is in contrast with suggestions that fast temporal relationships must be learned prior to slower ones in the motor (Krakauer et al., 2019) and language (Saffran & Wilson, 2003) learning literatures. It is possible that previous findings may reflect a bias in the strength of (simultaneous) learning for statistics that span different temporal scales, rather than a system constrained to learn rules at different timescales in a strictly sequential step-wise fashion. The lack of a strong correlation between human low and high-level sensitivity in the online prediction task is also consistent with this. Future work will have to confirm differences in the learning rate for statistical learning of input that spans different timescales, and confirm whether information presented at different timescales embedded in a single motor-perceptual stream is truly acquired simultaneously in humans. In addition, more work will be needed to tease apart the role of timescale per se vs. conceptual complexity and level of abstraction in explaining variance in learning trajectories for different temporal scales.

Our modeling results speak to how statistical dependencies at multiple timescales may be learned in the brain. We have previously developed a model of the hippocampus that provides an account of its role in rapid statistical learning (Schapiro et al., 2017). In the model, the monosynaptic pathway to region CA1 acts as a neural network with a single hidden layer, employing distributed representations and a relatively fast learning rate that allow it to effectively learn short timescale statistics quickly. Our current modeling results suggest that such a single-hidden-layer system may be able to concurrently handle longer timescale statistics. Temporal dependencies are known to be encoded at multiple timescales in both neocortex (Lerner et al., 2011; Hasson et al., 2015; Baldassano et al., 2017) and the hippocampus (reviewed in Davachi & DuBrow, 2015). In both cases, there appears to be an anatomical gradient of sensitivity to different timescales in different areas, with the hippocampus exhibiting increasing sensitivity to long timescales moving more anteriorly/ventrally along its long axis (Brunec et al., 2018; Raut et al., 2020; Bouffard et al., 2023). It may be that recurrent machinery allowing sensitivity to longer timescale statistics is increasingly present in more anterior/ventral segments of the hippocampus and/or its inputs. Neocortical gradients of timescale dependency may emerge for longer term forms of temporal knowledge.

In conclusion, humans are able to learn statistical information at multiple timescales within a short period, and their behavior can be effectively modeled using recurrent neural networks. Rather than learning rapid timescale statistics as a prerequisite for learning slower statistics, our model suggests that statistical information can be learned across multiple timescales simultaneously and via a shared mechanism and substrate. Although we may acquire rapid temporal regularities more readily than slowly evolving ones, the work demonstrates that learning one is not always a strong prerequisite for or barrier against learning the other.

## Acknowledgements

The authors thank Joey Zhao for assistance with data collection, and Elisabeth Karuza for helpful discussions.

## References

Baldassano, C., Chen, J., Zadbood, A., Pillow, J. W., Hasson, U., & Norman, K. A. (2017). Discovering event structure in continuous narrative perception and memory. Neuron, 95(3), 709–721.

Benjamini, Y., & Hochberg, Y. (1995). Controlling the False Discovery Rate: A Practical and Powerful Approach to Multiple Testing. Journal of the Royal Statistical Society: Series B (Methodological), 57(1), 289–300. 10.1111/j.2517-6161.1995.tb02031.x

Botvinick, M. M. (2007). Multilevel structure in behaviour and in the brain: A model of Fuster’s hierarchy. Philosophical Transactions of the Royal Society B: Biological Sciences, 362(1485), 1615–1626. 10.1098/rstb.2007.2056

Botvinick, M., & Plaut, D. C. (2004). Doing without schema hierarchies: A recurrent connectionist approach to normal and impaired routine sequential action. Psychological Review, 111(2), 395.

Bouffard, N. R., Golestani, A., Brunec, I. K., Bellana, B., Park, J. Y., Barense, M. D., & Moscovitch, M. (2023). Single voxel autocorrelation uncovers gradients of temporal dynamics in the hippocampus and entorhinal cortex during rest and navigation. Cerebral Cortex, 33(6), 3265–3283.

Brunec, I. K., Bellana, B., Ozubko, J. D., Man, V., Robin, J., Liu, Z.-X., Grady, C., Rosenbaum, R. S., Winocur, G., & Barense, M. D. (2018). Multiple scales of representation along the hippocampal anteroposterior axis in humans. Current Biology, 28(13), 2129–2135.

Chien, H.-Y. S., & Honey, C. J. (2020). Constructing and forgetting temporal context in the human cerebral cortex. Neuron, 106(4), 675–686.

Chung, J., Gulcehre, C., Cho, K., & Bengio, Y. (2014). Empirical evaluation of gated recurrent neural networks on sequence modeling. arXiv Preprint arXiv:1412.3555. https://arxiv.org/abs/1412.3555

Cleeremans, A., & McClelland, J. L. (1991). Learning the structure of event sequences. Journal of Experimental Psychology: General, 120(3), 235.

Creel, S. C., Newport, E. L., & Aslin, R. N. (2004). Distant melodies: Statistical learning of nonadjacent dependencies in tone sequences. Journal of Experimental Psychology: Learning, Memory, and Cognition, 30(5), 1119.

Davachi, L., & DuBrow, S. (2015). How the hippocampus preserves order: The role of prediction and context. Trends in Cognitive Sciences, 19(2), 92–99.

Fiser, J., & Aslin, R. N. (2002). Statistical learning of higher-order temporal structure from visual shape sequences. Journal of Experimental Psychology: Learning, Memory, and Cognition, 28(3), 458.

Friston, K. (2005). A theory of cortical responses. Philosophical Transactions of the Royal Society of London B: Biological Sciences, 360(1456), 815–836.

Furl, N., Kumar, S., Alter, K., Durrant, S., Shawe-Taylor, J., & Griffiths, T. D. (2011). Neural prediction of higher-order auditory sequence statistics. Neuroimage, 54(3), 2267–2277.

Glorot, X., & Bengio, Y. (2010). Understanding the difficulty of training deep feedforward neural networks. Proceedings of the Thirteenth International Conference on Artificial Intelligence and Statistics, 249–256. http://proceedings.mlr.press/v9/glorot10a

Gómez, R. L. (2002). Variability and Detection of Invariant Structure. Psychological Science, 13(5), 431–436. 10.1111/1467-9280.00476

Hasson, U., Chen, J., & Honey, C. J. (2015). Hierarchical process memory: Memory as an integral component of information processing. Trends in Cognitive Sciences, 19(6), 304–313.

Karuza, E. A., Kahn, A. E., Thompson-Schill, S. L., & Bassett, D. S. (2017). Process reveals structure: How a network is traversed mediates expectations about its architecture. Scientific Reports, 7(1), 12733.

Kiebel, S. J., Daunizeau, J., & Friston, K. J. (2008). A hierarchy of time-scales and the brain. PLoS Computational Biology, 4(11), e1000209.

Krakauer, J. W., Hadjiosif, A. M., Xu, J., Wong, A. L., & Haith, A. M. (2019). Motor learning. Compr Physiol, 9(2), 613–663.

Lerner, Y., Honey, C. J., Silbert, L. J., & Hasson, U. (2011). Topographic mapping of a hierarchy of temporal receptive windows using a narrated story. Journal of Neuroscience, 31(8), 2906–2915.

Lewicki, P., Czyzewska, M., & Hoffman, H. (1987). Unconscious acquisition of complex procedural knowledge. Journal of Experimental Psychology: Learning, Memory, and Cognition, 13(4), 523.

Misyak, J. B., Christiansen, M. H., & Tomblin, J. B. (2010). On-line individual differences in statistical learning predict language processing. Frontiers in Psychology, 1, 31.

Newport, E. L., & Aslin, R. N. (2004). Learning at a distance I. Statistical learning of non-adjacent dependencies. Cognitive Psychology, 48(2), 127–162.

Raut, R. V., Snyder, A. Z., & Raichle, M. E. (2020). Hierarchical dynamics as a macroscopic organizing principle of the human brain. Proceedings of the National Academy of Sciences, 117(34), 20890–20897. 10.1073/pnas.2003383117

Saffran, J. R., Aslin, R. N., & Newport, E. L. (1996). Statistical learning by 8-month-old infants. Science, 274(5294), 1926–1928.

Saffran, J. R., & Wilson, D. P. (2003). From Syllables to Syntax: Multilevel Statistical Learning by 12-Month-Old Infants. Infancy, 4(2), 273–284. 10.1207/S15327078IN0402_07

Sakai, K., Kitaguchi, K., & Hikosaka, O. (2003). Chunking during human visuomotor sequence learning. Experimental brain research, 152, 229–242.

Schapiro, A. C., Rogers, T. T., Cordova, N. I., Turk-Browne, N. B., & Botvinick, M. M. (2013). Neural representations of events arise from temporal community structure. Nature Neuroscience, 16(4), 486.

Shewalkar, A., Nyavanandi, D., & Ludwig, S. A. (2019). Performance Evaluation of Deep Neural Networks Applied to Speech Recognition: RNN, LSTM and GRU. Journal of Artificial Intelligence and Soft Computing Research, 9(4), 235–245. 10.2478/jaiscr-2019-0006

Shin, Y. S., & DuBrow, S. (2021). Structuring Memory Through Inference-Based Event Segmentation. Topics in Cognitive Science, 13(1), 106–127. 10.1111/tops.12505

